# A first perturbome of *Pseudomonas aeruginosa*: Identification of core genes related to multiple perturbations by a machine learning approach

**DOI:** 10.1101/2020.05.05.078477

**Authors:** Jose Arturo Molina Mora, Pablo Montero-Manso, Raquel García Batán, Rebeca Campos Sánchez, Jose Vilar Fernández, Fernando García Santamaría

## Abstract

Tolerance to stress conditions is vital for organismal survival, including bacteria under specific environmental conditions, antibiotics, and other perturbations. Some studies have described common modulation and shared genes during stress response to different types of disturbances (termed as perturbome), leading to the idea of central control at the molecular level. We implemented a robust machine learning approach to identify and describe genes associated with multiple perturbations or perturbome in a *Pseudomonas aeruginosa* PAO1 model.

Using microarray datasets from the Gene Expression Omnibus (GEO), we evaluated six approaches to rank and select genes: using two methodologies, data single partition (SP method) or multiple partitions (MP method) for training and testing datasets, we evaluated three classification algorithms (SVM Support Vector Machine, KNN K-Nearest neighbor and RF Random Forest). Gene expression patterns and topological features at the systems level were included to describe the perturbome elements.

We were able to select and describe 46 core response genes associated with multiple perturbations in *P. aeruginosa* PAO1 and it can be considered a first report of the *P. aeruginosa* perturbome. Molecular annotations, patterns in expression levels, and topological features in molecular networks revealed biological functions of biosynthesis, binding, and metabolism, many of them related to DNA damage repair and aerobic respiration in the context of tolerance to stress. We also discuss different issues related to implemented and assessed algorithms, including data partitioning, classification approaches, and metrics. Altogether, this work offers a different and robust framework to select genes using a machine learning approach.

## 1. Introduction

Cell stress can be defined as a wide range of molecular changes that cells undergo in response to environmental, physical, chemical, or biological stressors; sensing and responding to stress is critical for survival [1]. These biological functions and metabolic activities are executed through complex physical and regulatory interactions of genes that resemble networks [2]. Additionally, tolerance to stress conditions (i.e. stressors, perturbations, or disturbances) is vital for organismal survival, including bacteria under diverse environmental conditions, including antibiotics [1].

Several studies have revealed diverse molecular levels that can explain the general response to disturbances in many organisms. However, detailed mechanisms related to responses to perturbations remain poorly understood [3]. Available reports usually focus on specific stressors and relatively few studies have focused on common, central, and universal determinants affected by multiple perturbations [3]. This concept has been recently referred to as the perturbome [4,5]. For example, in eukaryotic organisms, including plants and human cancer models, some studies have described diverse stress-response genes as common modulators for different types of disturbances, suggesting a central control mechanism [2,4–6]. In prokaryotic models, similar findings have been reported for *Escherichia coli* [3,7].

Additionally, a comprehensive study of gene-interactions allows the identification of functional relationships among genes [8], their products, and the underlying biological phenomena that are critical to understand phenotypes under different biological conditions [9,10]. In the context of cell stress, the response to different environmental or experimental stimuli can be recognized by distinct gene expression patterns. This can be inferred from transcriptomic profiling data and functional associations using high throughput molecular technologies such as microarrays or RNASeq [2]. However, a challenge with these technologies is the large amount of high complexity data generated. Specialized bioinformatics analyses are required to select relevant information and to reduce noise that distinguishes the molecular determinants for particular biological conditions. Thus, a primary objective of transcriptomic profiling is to find an optimal subset of genes that could be used to characterize and classify unknown samples [11]. This gene selection is not obvious and complex due to the thousands of genes to select from [12].

To study the central response determinants to different perturbations in a living organism, we used the model *Pseudomonas aeruginosa* PAO1 (the reference strain of Pseudomonals). *P. aeruginosa* is a Gram-negative gamma-proteobacterium with a noteworthy metabolic versatility and adaptability enabling the colonization of diverse niches infecting plants, animals, and humans alike [8,13]. This bacterium is an opportunistic pathogen that causes a variety of infections in immunocompromised hosts, and it is associated with complications in cystic fibrosis and is a major cause of death among patients with this disease [14]. For Pseudomonals, the molecular mechanisms associated with most biological processes remain unclear causing limited action to modulate responses, including the susceptibility to stressors, environmental factors, and experimental conditions.

In *P. aeruginosa*, the response to disturbances has been reported to be influenced by <10% of coding genes [15,16], including complex molecular networks with pleiotropic patterns [17], i.e., a single perturbation induces the modulation of hundreds of genes [15,16]. These networks can be modeled using system biology approaches to recognized emergent patterns at the molecular level, as previously reported [18–21]. Large-scale molecular networks are regularly built using databases and can be used not only to understand biological functions, regulatory mechanisms, pathways, or gene groups associated with the molecular response [22] but also to eventually manipulate it [23].

In the response to stress conditions, gene expression profiling using transcriptomic analysis and topological metrics can be used to provide a global view of transcriptional regulation [24]. The identification of hub genes and pathways associated with the network can be used to gain insights into the biological meaning of the complex interactions [24,25]. This is a key point to translate molecular patterns into the phenotypes, in particular for the response to perturbations [26].

Several studies have used machine learning algorithms at the transcriptomic level to recognize gene expression patterns [2,27–29]. However, for many common biological contexts, the applicability and utility of these machine learning approaches have not been fully explored and utilized [30], for example in the exposure to multiple stressors and the molecular response. To our knowledge, only a few studies have used feature selection methods on biological data to describe the effects of multiple perturbations in complex biological systems [4,6] and so far none in *P. aeruginosa*. A related work in *P. aeruginosa* used a machine learning approach to identify sets of genes that correlate with multiple culture media [31]. Due to differences in the experimental design, this study and our work are not comparable.

The use of microarray and other high throughput technologies data involve challenges for machine learning approaches. These include the curse of dimensionality [32,33], normalization of raw values to compare samples [34,35], data partitions for training and testing models [36,37], and evaluation of performance [34,38]. Since comparison between the machine learning algorithms is completely variable [11,30,33,39,40], Support Vector Machine (SVM) [41], K-Nearest neighbor (KNN) [40], and Random Forest (RF) [42] have been successfully used with microarray gene expression data allowing the recognition of emerging patterns [39].

Here, we hypothesize that perturbations on living cells will trigger global reprogramming of multiple molecular determinants that can be sensed at the transcriptional level. The initial central response after acute stress will then expand producing the global molecular rearrangement (Figure 1-A). Then, pleiotropic and specific effects on gene networks will be reflected as changes in gene expression profiles and the complexity of molecular regulation at other levels. Therefore, by using a machine learning approach, common molecular features (for all stressors) could be identified as a central or core determinant.

**Figure 1.**
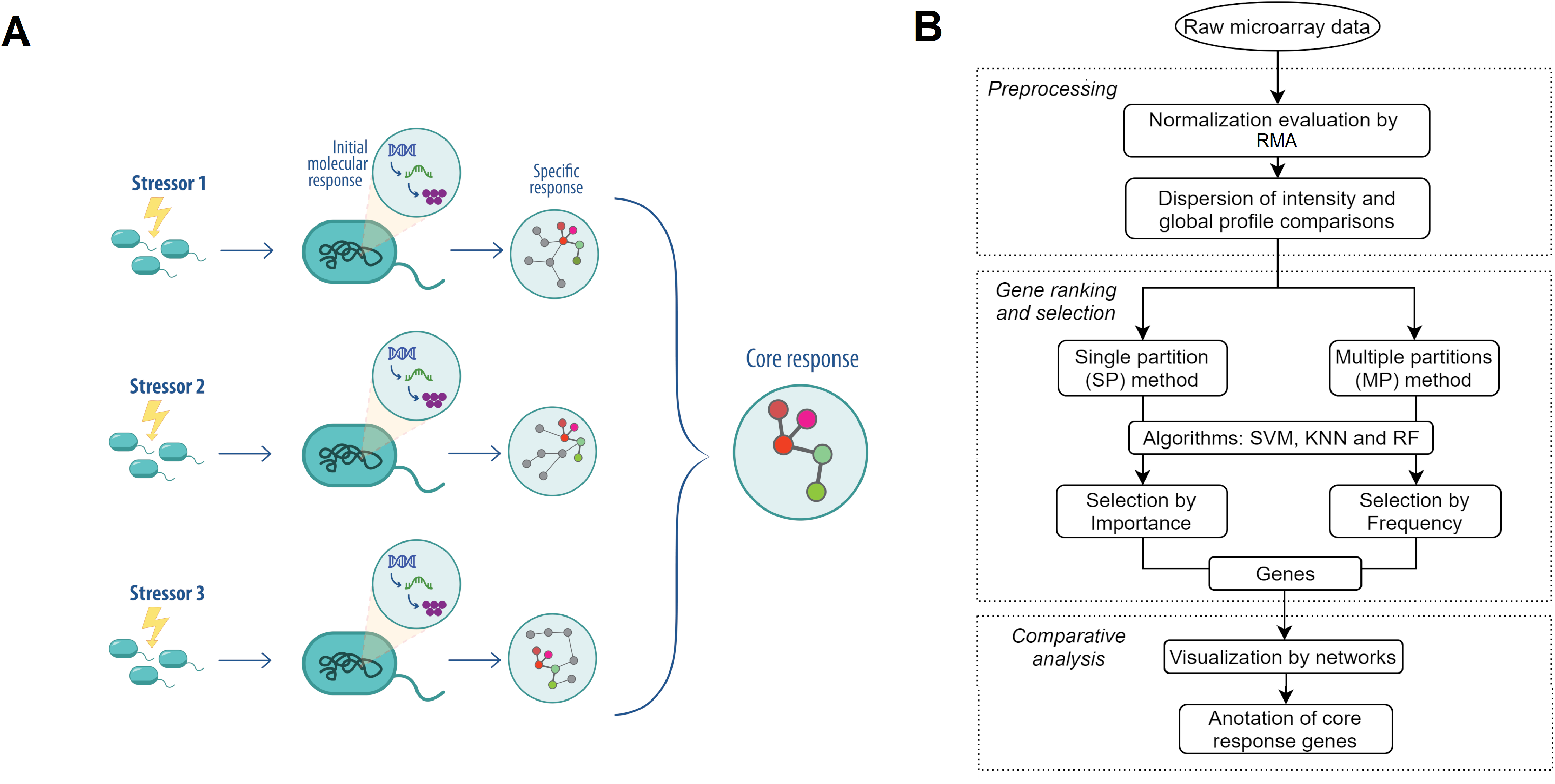
General pipeline for identifying core response genes in *P. aeruginosa* by a machine learning approach. (A) Schematic representation of hypothesis for identifying core response determinants when bacteria are exposed to multiple perturbations. (B) The workflow of the machine learning approach using microarray data and model fitting by SP and MP methods for identifying and describing core response genes.

Thus, the aims for this study were (i) to implement a machine learning approach to select genes from gene expression data, and (ii) to identify and describe a subset of genes that can be associated with multiple perturbations or the core perturbome in *P. aeruginosa.*

## 2. Materials and Methods

The overall methodology of the study is presented in Figure 1-B. In brief, after a data selection, normalization and gene selection to define the perturbome were run. To achieve this, we considered two different procedures to split the data (training and testing datasets for the machine learning approaches: SVM, KNN, and RF): a single partition (SP method) and another with multiple partitions (MP method). Relations between genes were represented using both large scale and small-world networks, and a final comparison between conditions, an analysis of differential expression, and gene annotation were performed.

### 2.1. Datasets

To compare gene expression profiles of strain *P. aeruginosa* PAO1 when exposed to multiple perturbations, the GEO database (https://www.ncbi.nlm.nih.gov/geo/) was used for a systematic selection of datasets (Figure 2-A-B). Initial evaluation by the organism and mRNA profiles by GPL84 platform (Affymetrix *Pseudomonas aeruginosa* PAO1 Array, with all 5549 protein-coding sequences) identified 156 series of datasets with 1310 samples (Date of Access: January 25th, 2018). In a second step, data were selected according to experimental design if they included perturbations, leaving only 47 series. Finally, to make datasets comparable by experimental conditions, evaluation and selection were done for series with similar culture conditions (Luria Bertani LB medium and exponential phase when measuring mRNA profile) and if a control condition was available (without any perturbation or treatment). The final dataset was composed of 10 series with 71 samples (Series GSE2430, GSE3090, GSE4152, GSE5443, GSE7402, GSE10605, GSE12738, GSE13252, GSE14253 and GSE36753).

**Figure 2.**
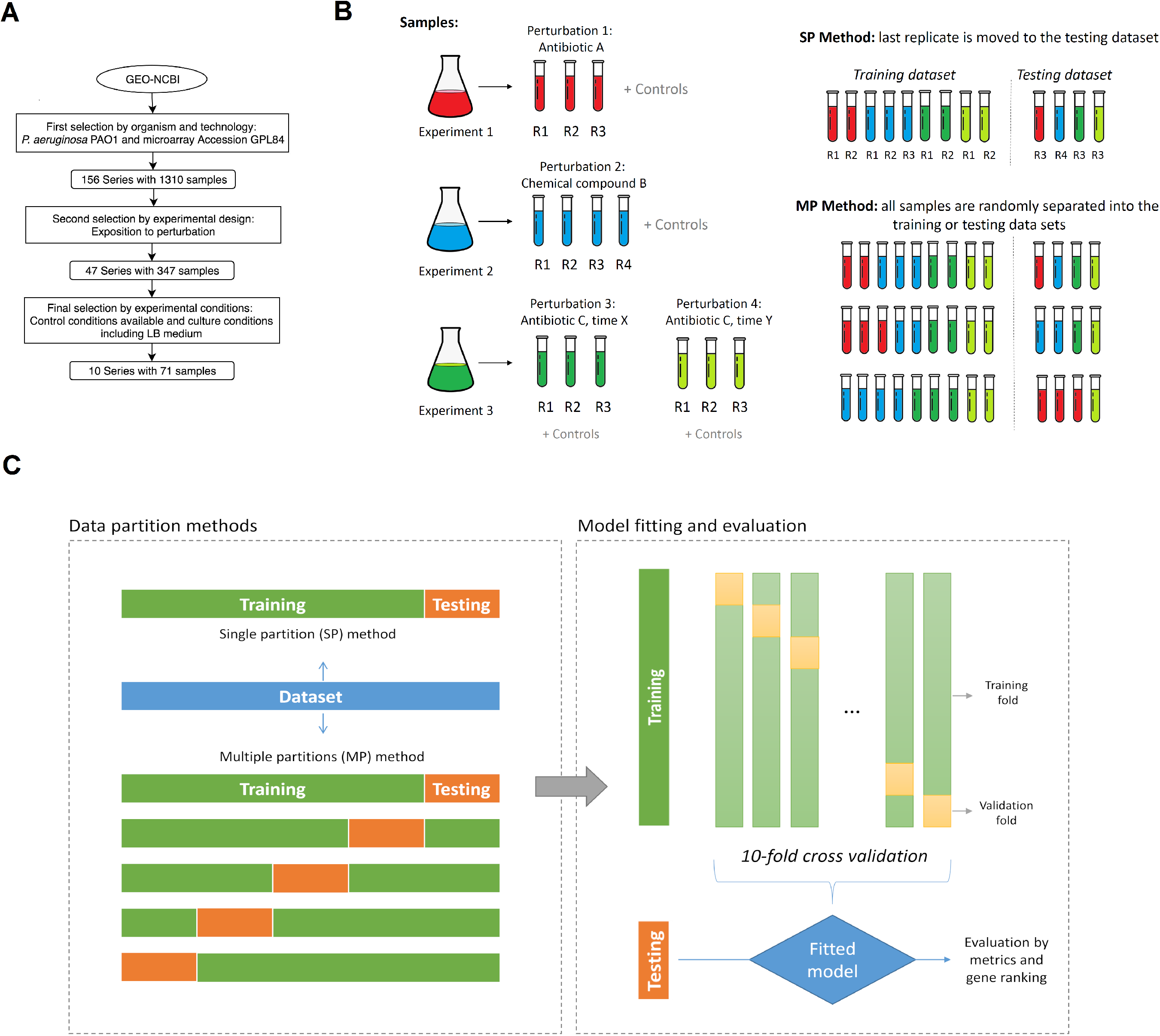
Data selection and partitioning. (A) Microarray-data selection in GEO-NCBI database according to experimental design. (B) Samples, including replicates in each experiment −perturbation and control-, were separated into the training or the testing data sets using the SP (the last replicate of each condition was moved to the testing dataset) or MP (random separation with multiple arrangements) methods. (C) Representation of classification procedures for the SP and MP methods, including internal 10-fold cross-validation for the training data set.

Some series included temporal measurements (see Experiment 3 of the Figure 2-B) which we considered as separate perturbations, resulting in replicates of 10 controls and 14 perturbations (Figure 3-A): azithromycin with 2 series (AZM-a and AZM-b) [43,44], Hydrogen peroxide (H2O2) [45], copper (Cu) [46], sodium hypoclorite (NaClO) [47], ortho-phenylphenol (OPP) at 20 and 60 minutes [48], colistin (COL) [49], chlorhexidine diacetate (CDA) at 10 and 60 minutes [50], E-4-bromo-5-bromomethylene-3-methylfuran-2-5H-one (BF8) [51] and ciprofloxacin CIP at 0, 30 and 120 minutes [19]. Details of each study (experimental design and main results) are provided in Table S1.

**Figure 3.**
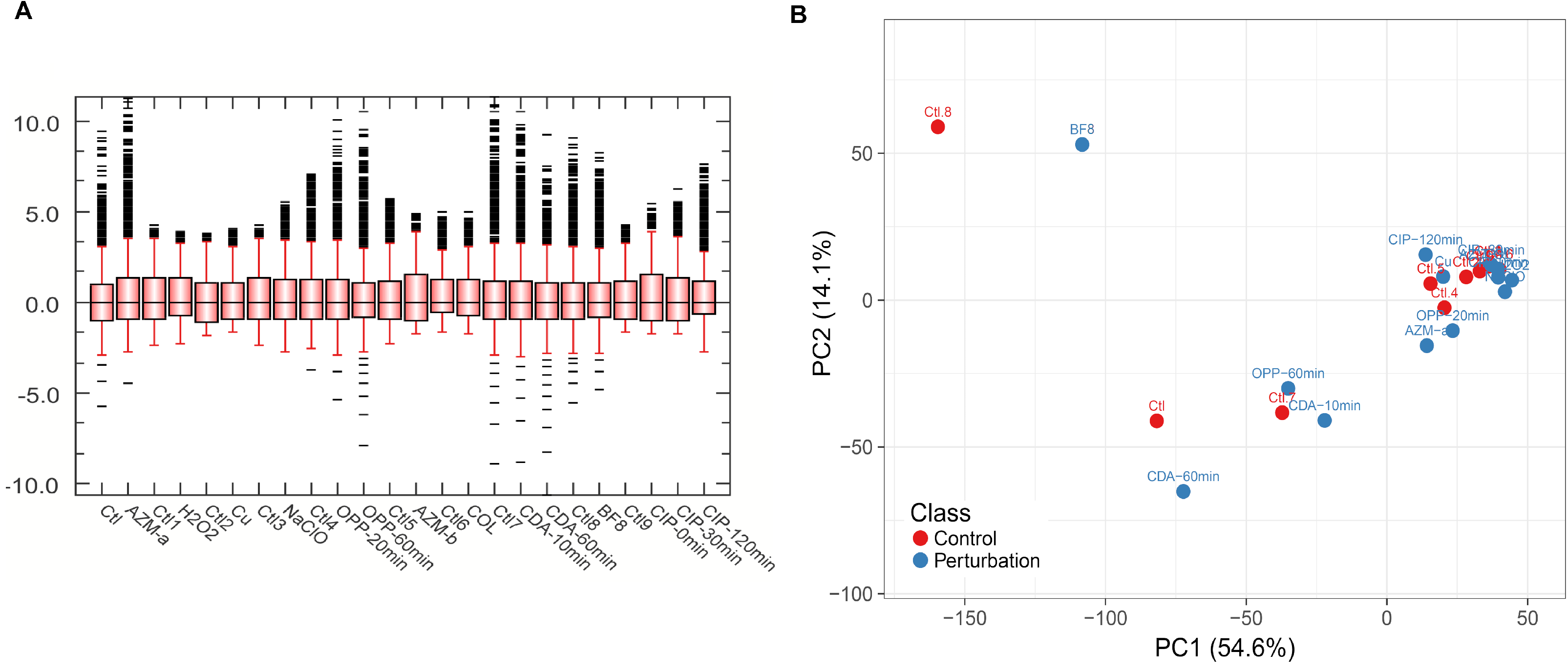
Comparison of samples by global profiles with transcriptomic data of 5549 genes of *P. aeruginosa*. (A) Dispersion of intensities of samples, showing similar distribution between samples after normalization. (B) Global profiles of samples were compared by PCA, using all the 5549 genes in the microarray.

### 2.2. Pre-processing and comparison of global transcriptomic profiles

To compare all the 71 samples, a first analysis was the pre-processing step using Bioconductor 3.8 (https://www.bioconductor.org/) in the R software (Version 3.5) with classical functions for microarrays. Robust MultiArray Average algorithm (RMA) was performed in the Affy package to correct background, normalization, and summarization.

Subsequently, a clustering algorithm of Principal Component Analysis (PCA) was implemented to compare global transcriptomic profiles between perturbations and controls. PCA was done with default parameters of the Caret Package (caret.r-forge.r-project.org/) in R software.

In order to robustly select genes that could separate experimental conditions (controls and perturbations) and to identify the core response of *P. aeruginosa*, two approaches of feature selection protocols were implemented, as detailed below (Figure 2-B-C).

### 2.3. Gene ranking and selection by Single Partition SP method

To identify genes which expression values were commonly related to multiple perturbations, a first approach was implemented considering a particular partition of the dataset for training and testing sets (Figure 2-B-C). Single partition was established using the last replicate of each experiment, in both control and perturbation. Because there were 14 perturbations and 10 controls (71 samples including replicates or individual samples), a total of 24 samples were included in the testing dataset and the remaining 47 samples were included for the training dataset (66% for training and 34% for the testing set).

Using this partition, the ranking of genes was done by a machine learning approach using three classification algorithms: SVM, KNN, and RF. A homemade script in R included these functions of the Caret package. For all three algorithms, default parameters were used for training and 10-fold cross-validation was included. After this, the variable importance metric was calculated for all genes using the *varImp* function, associating a specific value for each gene. In the case of SVM and KNN, the same importance is calculated because the function is model-free for these cases (as detailed in Caret Package), resulting in the same list of genes but metrics are specifically calculated for each algorithm. For RF, the accuracy and model parameters were used to calculate the metric.

As similarly reported in [11], to evaluate the number of genes that should be selected in the top group (the first K ranked genes, as candidates for the core response by each algorithm), multiple classification models were systematically run, starting only with the highest-score ranked gene. In brief, after the training with one gene, model performance was evaluated by calculating metrics using the testing dataset. Then, training was run again when the next ranked gene was added, and new metrics were calculated. This process was repeated up to complete all the ranked genes. Metrics included accuracy (correct classification percentage), kappa value (inter-rater classification agreement), sensitivity, specificity, precision, recall, prevalence, and F1 score (harmonic average of the precision and recall).

Selection of the K value of top genes was based on the following criteria: (i) the stability of the metrics (priority for accuracy, kappa, and F1) when the increment of ranked genes was done, as suggested in [11], (ii) consensus value suitable for all the three algorithms (including a 10% of tolerance), and (iii) minimum number of genes as possible. After the selection of the K value, ROC (Receiver-operating characteristic) curve and AUC (Area under the curve) value were calculated for each algorithm. Finally, the selection of top K genes between algorithms was compared by metrics and- the list of genes.

### 2.4. Gene ranking and selection by Multiple Partitions (MP) method

In order to identify genes related to multiple perturbations independently of a single/specific partition, a second method using multiple random partitions was implemented (Figure 2-B-C). To address this, a random data selection for training and testing sets was done using the *createDataPartition* function. Partition was set to 80% (57 samples) for the training set and the remaining for the testing set (14 samples) with experimental conditions equally distributed. Then, protocols with SVM, KNN, and RF algorithms (same conditions as previously described in SP method, with 10-fold cross-validation) resumed the analysis with a final ranking of genes using the *varImp* function. Using only the top K of ranked genes (K value determined using the criteria described in the SP method), a new set of training/testing sets were used for evaluating the performance of the models and each metric was stored with the list of the K ranked genes. This full process was automatically repeated 100 times using the *replicate* function, starting with a new random partition and finishing with the list of the K genes and the metrics associated with that partition. Finally, for each algorithm, full data of all the runs were analyzed for determining the frequency of the appearance of genes (*table* function) and calculating the average and dispersion of metrics across all the 100 runs. The definitive list of the K more frequent genes was established for each algorithm after this comparison by frequency.

### 2.5. Identification of core response genes

After selection of top K genes (candidates of the perturbome) in each algorithm by SP and MP methods, a comparison of genes was done using UpSets plots (https://github.com/hms-dbmi/UpSetR) in R software. We called the “support” of a gene the number of algorithms (out of six approaches) in which a gene was selected as a candidate. Genes with a support ≥4 (i.e., it was identified by at least four algorithms) were considered as part of the final list of genes of the core perturbome. This consideration guaranteed that a gene was identified by the two methods and at least by two different classification algorithms.

Gene relationships were represented using molecular networks using a large scale model (using a top-down systems biology approach). The network was built using protein-protein interaction (PPI) graphs in PseudomonasNet database (www.inetbio.org/pseudomonasnet/). Visualization of the network and measurement of the topological features were performed using Cytoscape software [52].

### 2.6. Description and comparison of core response genes

In order to describe and compare the genes associated with the core response of *P. aeruginosa*, by experimental conditions, four analyzes were established. First, PCA was evaluated again but now only considering genes of the core response. Based on distribution in the case of PCA, representation of centroids was done using the Kmeans algorithm.

Second, using the PseudomonasNet database, a small-world network was built and then exported into Cytoscape software with the genes of the core response. The information of topological features (including connectivity) and expression levels of kmedoids were incorporated into different versions of the network. To identify central or hub genes in the network in Cytoscape, the cytoHubba app was used [53] with the top 5 elements according to the metrics of degree, bottleneck, and betweenness. All identified genes in any out of the three approaches were assigned as hub genes.

Third, a classic analysis of differentially expressed genes (DEGs, *p<0.05)* was implemented in R with Limma package (https://www.rdocumentation.org/packages/limma/versions/3.28.14) using empirical Bayes moderated t-statistic (eBayes) with Benjamini and Hochberg method for *p*-value correction [54]. This was done to compare our results with a classical approach for gene expression.

Finally, to give biological interpretation to the selected genes, to determine levels of expression reported in databases, and to study metabolic pathways involved under each perturbation, an exhaustive annotation was made using the databases PseudomonasDB (gene ontology), GEO database, KEGG annotation, and particular literature. This information was integrated with the results obtained by all the analysis and the DEGs to fully describe the genes that make up the core response or perturbome of *P. aeruginosa* PAO1.

#### Availability of data and material

The datasets generated and/or analyzed during the current study are available in:

- Public raw data used in this study: GEO database (https://www.ncbi.nlm.nih.gov/geo/), data Series GSE2430, GSE3090, GSE4152, GSE5443, GSE7402, GSE10605, GSE12738, GSE13252, GSE14253 and GSE36753.
- Normalized data and R Scripts: https://github.com/josemolina6/CoreResponsePae

## 3. Results

### 3.1. *Perturbome genes of* P. aeruginosa *can be identified by a machine learning approach using SP and MP methods*

A total of 71 samples of 10 controls and 14 perturbations were considered for the study, with comparable expression levels (Figure 3-A). Global transcriptomic profiles (all 5549 genes) were compared by a PCA, revealing a mixed pattern (no separation) between perturbations and controls (Figure 3-B). Two samples (BF8 and Control 8) resulted in “extreme” global profiles. Two methods using machine learning (SP and MP) were implemented in order to robustly rank and select genes associated with multiple perturbations in *P. aeruginosa* (Figure 2). In each method, three classification algorithms were evaluated: SVM, KNN, and RF.

The first method (SP) implemented a gene ranking based on variable importance using a single/specific data partition. Metrics associated with RF are shown in Figure 4, and in supplementary Figure S1 for SVM and KNN. After the ranking was established, multiple classification models were run with the ranked genes (Figures 4-A and supplementary Figure S1-A-C). For each classification model, the stability of the three metrics was evaluated to select the suitable K value of genes that could be applied to all algorithms at the same time. For SVM and RF, stable values of metrics are given with at least the first 51 ranked genes, meanwhile it is 45 genes for KNN. Considering a 10% of tolerance with the highest of these values, K=56 was selected as the number of top genes that were included as preliminary candidates of the core response according to each algorithm. With this value, metrics of each algorithm were compared (Table S2). For example, accuracy was 0.79, 0.71, and 0.75 for SVM, KNN, and RF respectively in the SP method. SVM obtained a better performance according to kappa, sensitivity, recall, and F1 scores, but higher values of specificity and precision resulted for RF. Also, the ROC curve and AUC value were calculated (Figures 4-B and supplementary Figure S1-B-D). The best performance was obtained for RF with AUC = 0.92, then 0.82 for SVM, and finally 0.76 for KNN. Since the ranking by *importance* for SVM and KNN is the same (model-free strategy, see details in https://topepo.github.io/caret/variable-importance.html), they shared the same list of top 56 genes. Comparison between implementations showed that 21 genes were identified by the three algorithms at the same time, 35 exclusively by RF, and the same number for SVM/KNN. In total 91 genes were identified by any of the algorithms. The list of genes and the importance value of the top 56 ranked genes for each approach is presented in Figures 4-C for RF and Figure S1-E for SVM/KNN.

**Figure 4.**
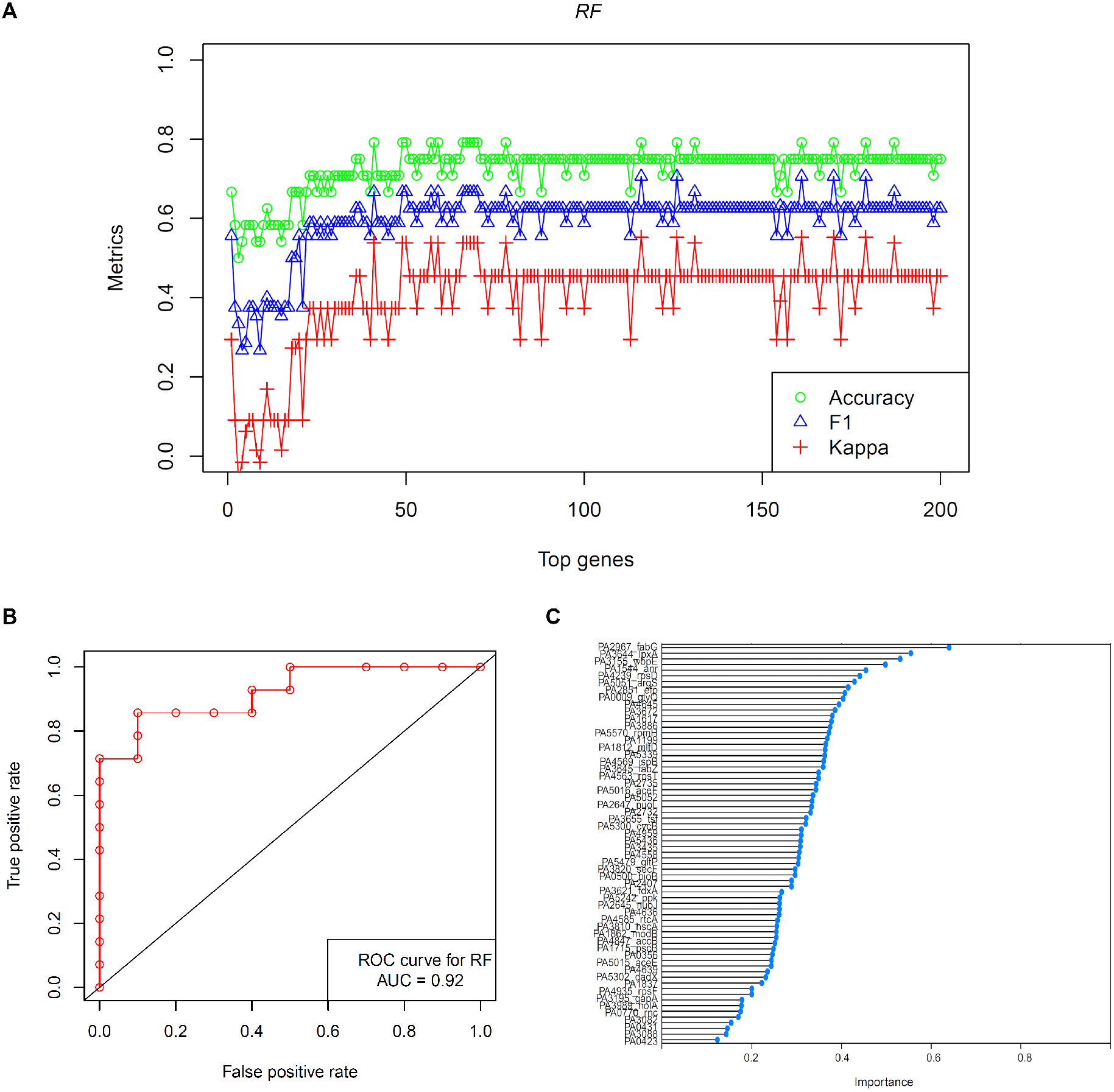
Evaluation of SP method for gene ranking by importance, the case for RF algorithm. (A) Accuracy, F1, and kappa values after iterations of classification with the first top 200 genes (adding genes 1-to-1). (B) ROC plot using selected top 56 genes for evaluation of the performance of the algorithm. (C) Ranking and importance value of top 56 genes. Similar results are shown for SVM and KNN algorithms in supplementary Figure S1.

In a second approach, the MP method was implemented using multiple random partitions. Same SVM, KNN, and RF algorithms were evaluated by running 100 iterations with different partitions and top-56 more frequent genes for each method were selected. Details of ranking and frequency are shown in Figure 5-A (SVM) and supplementary Figure S2 (KNN and RF). Metrics for all 100 iterations are presented in Table S2, Figure 5-B, and supplementary Figure S2. Accuracies for all the models were 0.66, 0.69, and 0.70 for SVM, KNN, and RF respectively. Specific values for kappa, precision, recall, and F1 score were found for each algorithm. When a comparison of the list of genes was done, 29 genes were identified at the same time by all implementations, and 23 by both SVM and KNN.

**Figure 5.**
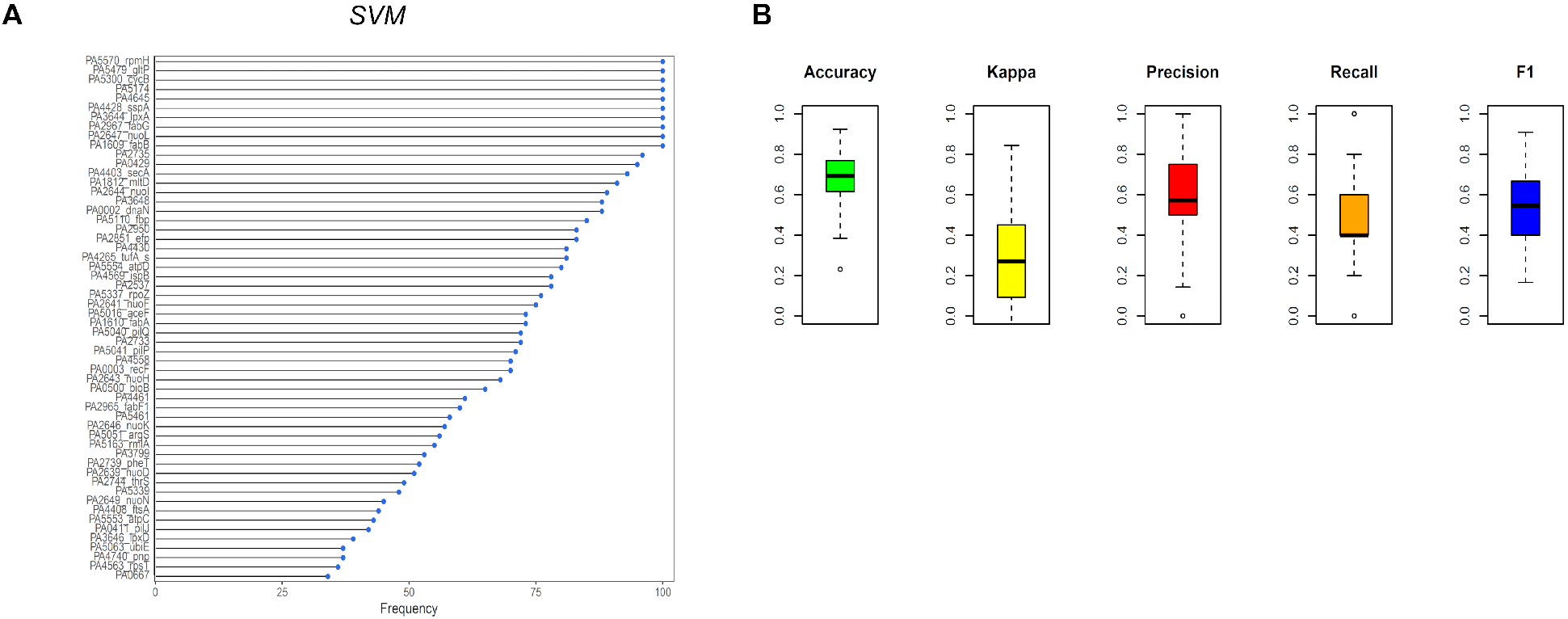
Evaluation of MP method for gene ranking by frequency, case for SVM algorithm. (A) Ranking of top 56 genes by frequency after iterations of 100 data partitions and classification model fitting. (B) Dispersion of metrics across 100 iterations. Similar results are shown for KNN and RF algorithms in supplementary Figure S2.

To identify preliminary core response genes in *P. aeruginosa*, the lists of top 56 genes selected by each algorithm and method were jointly represented using UpSet plots (Figure 6-A). A total of 118 different genes were identified and 15 genes were simultaneously identified by all the algorithms. The KEGG annotation of all the genes revealed functions associated with energy and DNA metabolisms, biosynthesis, ribosomal activity, and other pathways (Figure 6-B). See below for more details of the annotation.

**Figure 6.**
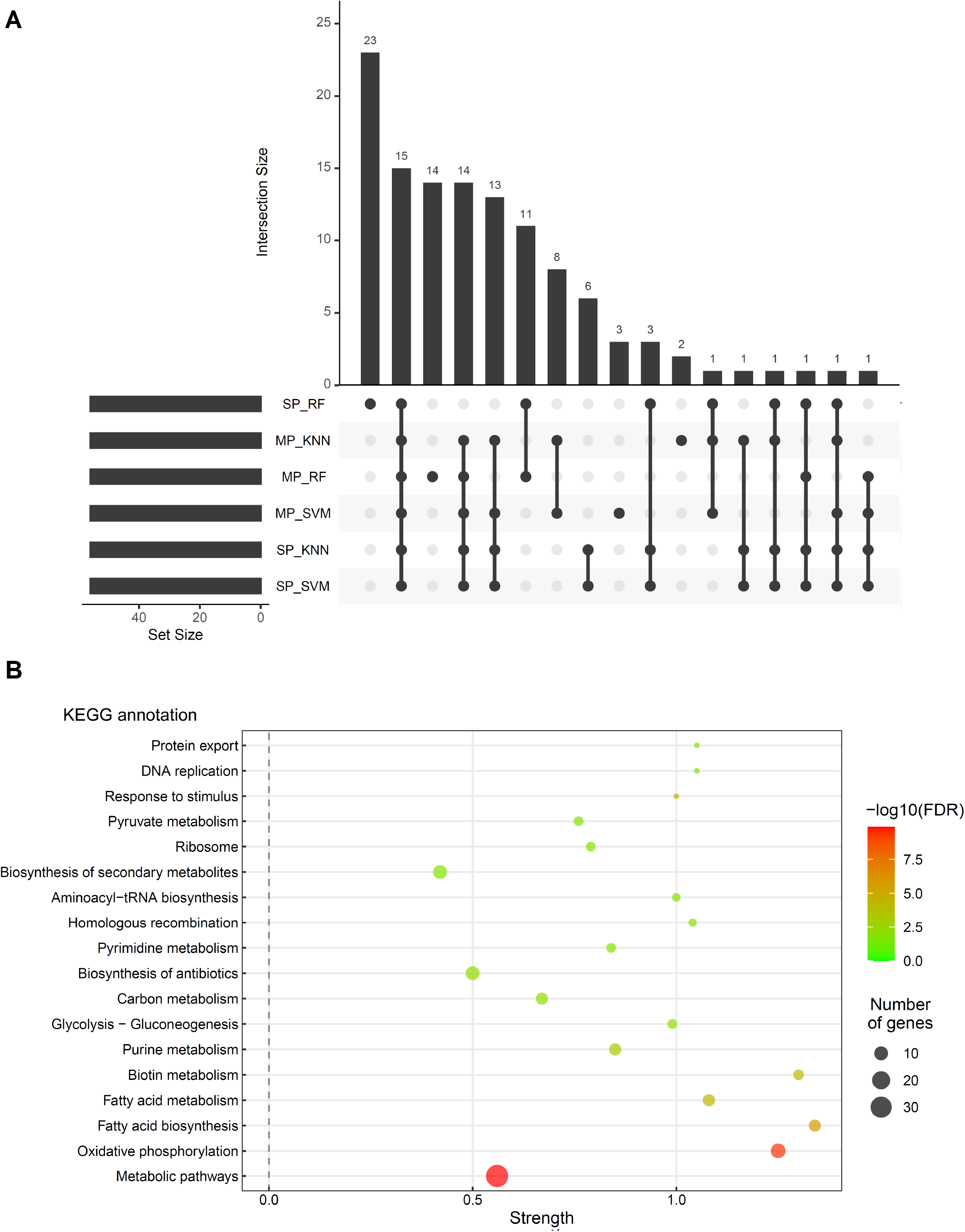
Comparison of 118 candidate genes of the core perturbome. (A) The top 56 genes of each algorithm were selected and compared using upset plots, showing the intersection of the lists among classifiers. This comparison resulted in 118 distinct candidate genes. A gene was selected in the final list if it was found at least by four different algorithms, i.e. the support was ≥ 4. (B) Gene annotation using KEGG shows the enriched pathways found for the core perturbome genes.

The distribution of all 118 genes on a large-scale molecular network is presented in Figure 7-A. Results show that selected genes are associated with different subgroups of highly connected genes.

**Figure 7.**
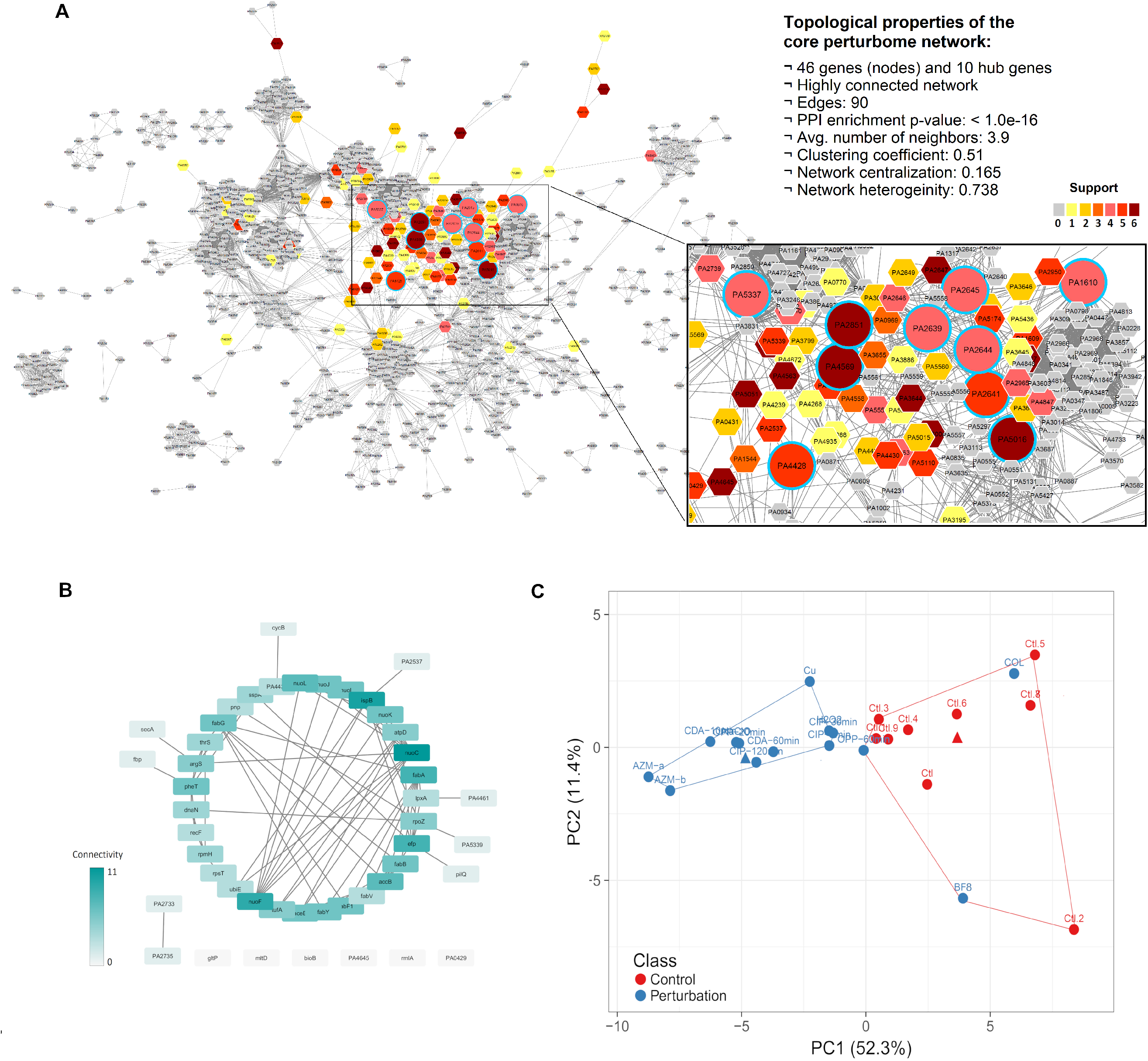
Network analysis and comparison of global transcriptomic profiles using the 46 core perturbome genes. (A) Distribution and support (number of algorithms) of preliminary 118 selected genes on a basal large-scale network of functional associations in *P. aeruginosa.* Topological metrics of the network and identity of hub genes (circles with light-blue border) are included. (B) Small-world network showing relationships between the 46 core response genes and connectivity metric for each gene. (C) The global profile of samples by PCA, showing the separation of conditions. For reference, centroids were plotted as triangles in each cluster. More details are shown in Supplementary Figure S3.

The topological metrics of the network showed a non-random arrangement with 90 edges (more than expected by chance, enrichment p-value: < 1.0e-16), 3.9 average number of neighbors, a clustering coefficient of 0.51, and others. Besides, 10 hub genes were identified among the core perturbome, which are representative elements of the network. More details in Figure 7-A and Table 1.

**Table 1.**
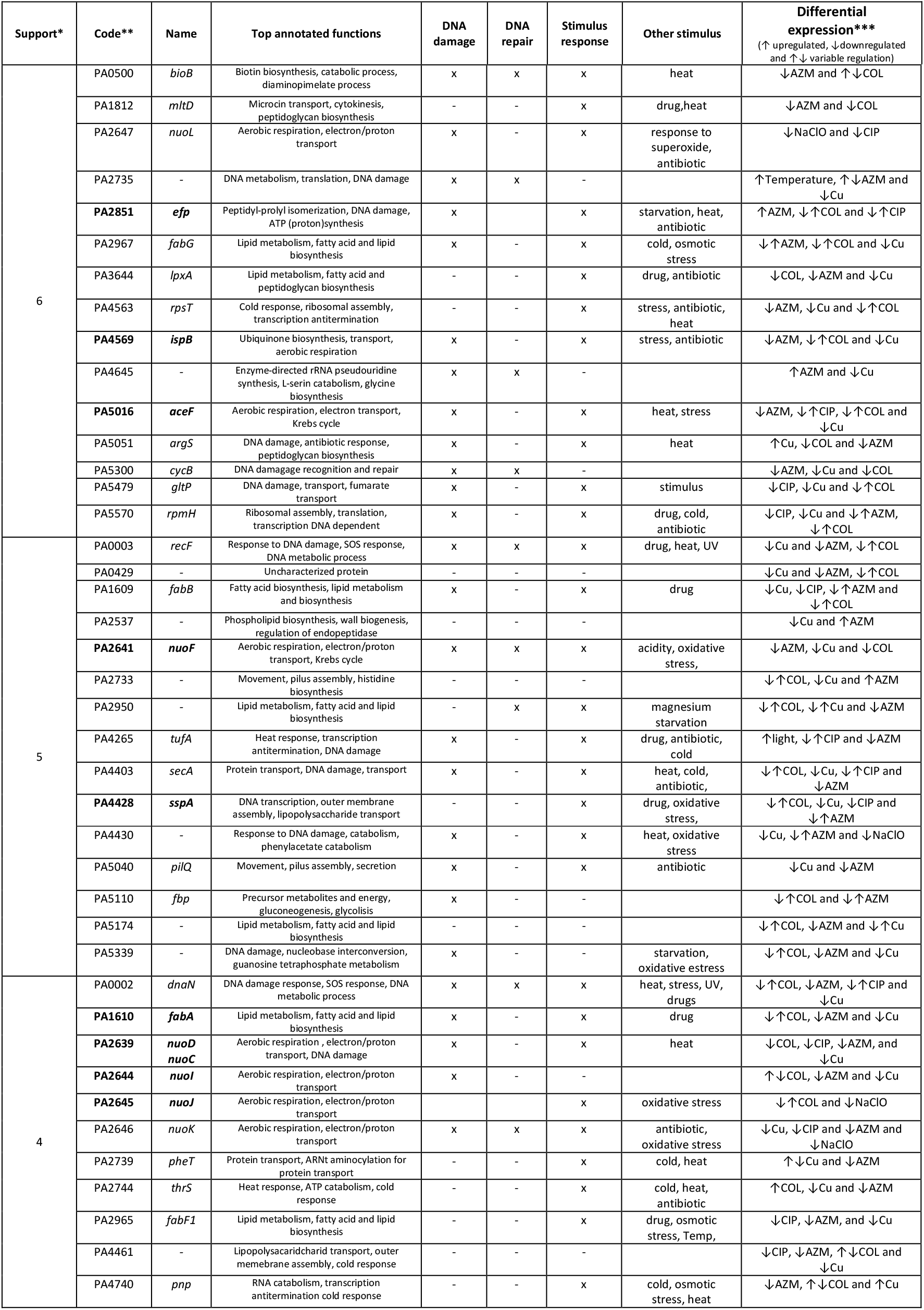
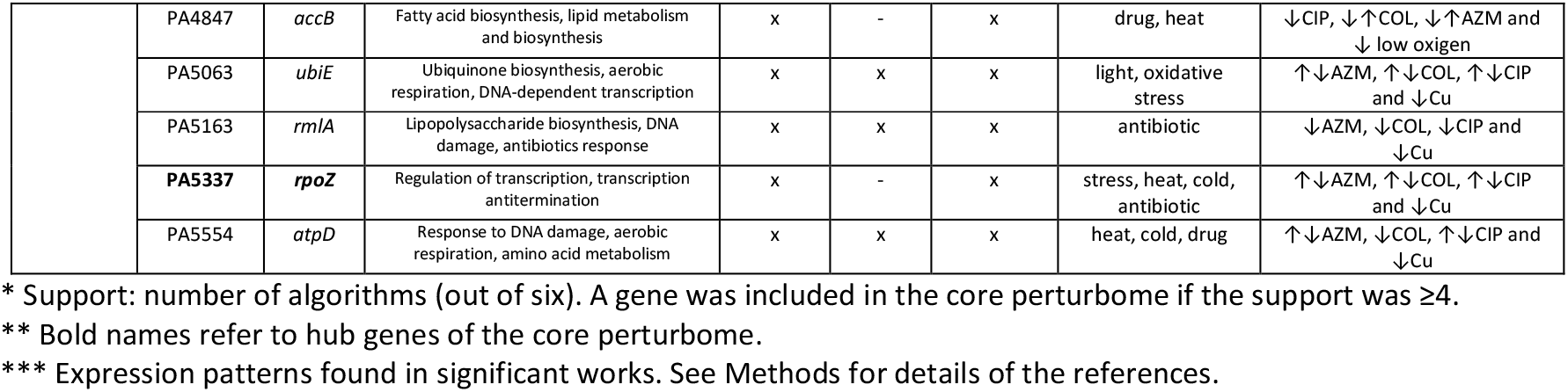
Annotation of genes of the core perturbome *Pseudomonas aeruginosa*

The final version of the core perturbome was established by selecting genes with a support ≥4 (i.e found by at least four algorithms). In total 46 genes were finally associated with core response. Notwithstanding six individual classifiers were implemented (each one with specific datasets), the consensus strategy to select the final genes (by the support) had no effects on the resolution of the classes (control or perturbation), as shown with the PCA with the transcriptomic profile using the 46 genes.

In the representation as a small-world network, relationships between the 46 genes revealed different topological patterns of connectivity between molecules, being *nuoC* and *nuoF* the ones with higher connectivity (connection degree). In addition, only six genes had no connections between them (Figure 7-B).

### 3.2. Comparisons of core response genes show the separation of global transcriptomic profiles according to experimental conditions and biological functions related to tolerance to stress

After the selection of the core response genes, comparisons between controls and perturbations were done to characterize these genes by global profiles. First, the PCA was implemented to compare the global profiles using the 46 genes of the perturbome, in which the distribution shows a well-defined separation between controls and perturbations (Figure 7-C). A Kmedoid algorithm (k = 2) was able to identify two clusters enriched by samples of each condition: one consisting entirely of perturbation samples (11 samples, blue color) and the other mostly by control samples (10 controls and 3 disturbances, red color). In the case of the blue cluster, the kmedoid was sample CIP-120min, meanwhile control 6 was selected for another cluster. Supplementary analysis of gene expression was included by comparing levels for all core response genes (Figure S3-A) and comparing expression levels of the kmedoids on the small-world network (Figure S3-B-C).

On the other hand, gene annotation revealed that biological processes related to most of the genes are metabolism, molecule binding, and biosynthesis (Figure 8-A). At the level of pathways, genes of the core perturbome have participation in processes of DNA damage response, DNA repairing, and response to general/specific stimuli (Figure 6-B and Table 1). Other studies show distinct expression patterns for each gene, depending on the disturbance as shown in Table 1. Finally, in order to compare the results of the machine learning strategy with another approach, differential expression analysis was run. A total of 101 DEGs were identified, of which 33 were shared by the core response (Figure 8-B). This means that 72% of core response genes were also identified by another single and independent method. Annotation of DEGs for biological processes (Figure 8-C) showed a similar profile to our machine learning approach.

**Figure 8.**
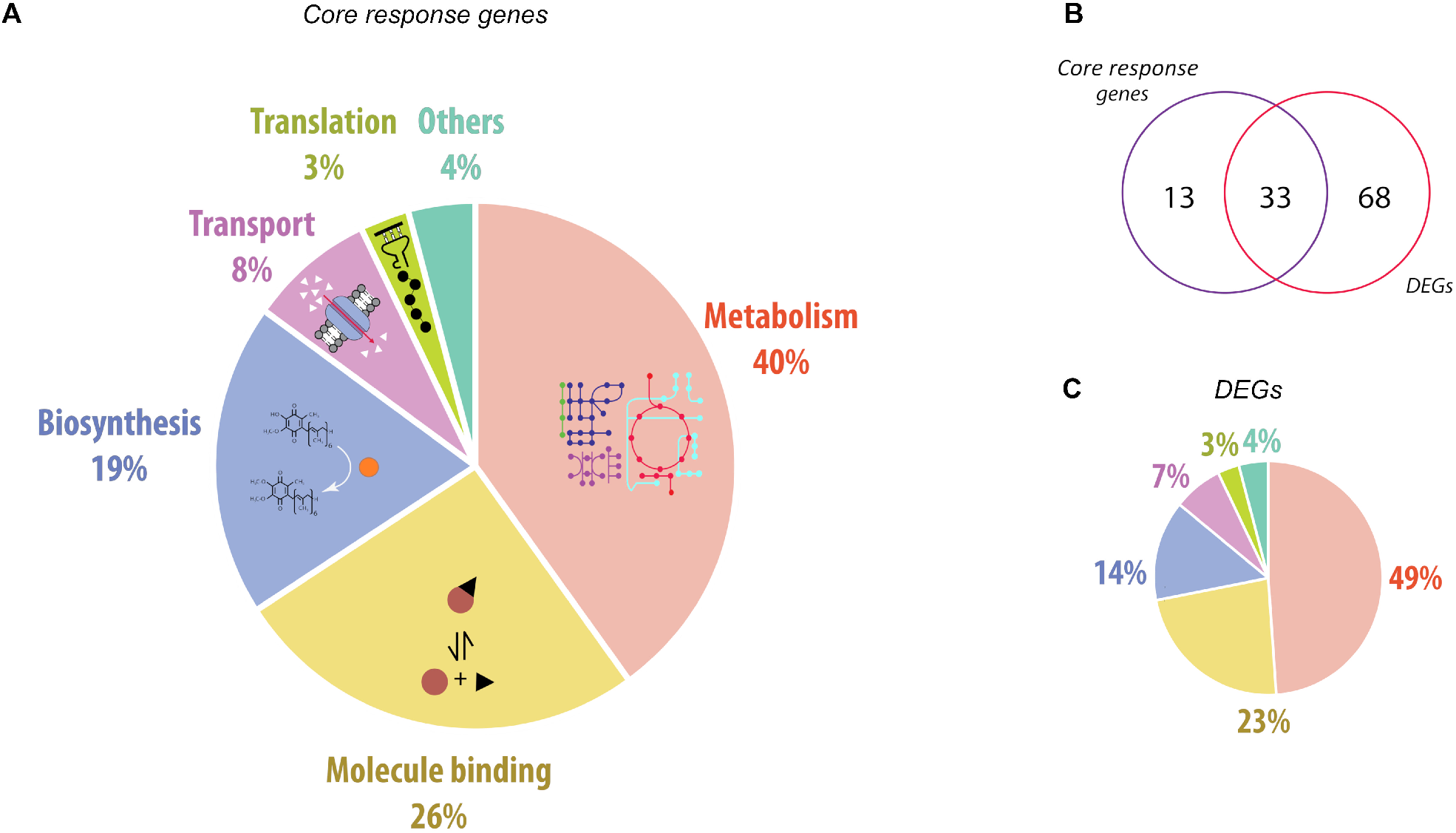
Annotation of core response genes and comparison with a differential expression analysis. (A) General annotation profile of identified genes by core response genes showing associated biological processes. (B) Comparison of identified genes by our machine learning approach and DEGs lists. (C) General annotation of DEGs showing a similar profile to our approach.

## 4. Discussion

Living organisms face external and internal conditions that compromise cellular functions at molecular, metabolic, and structural levels, disrupting their homeostasis [55,56]. The cell stress response is crucial for organismal survival and complex networks are usually involved in the molecular mechanism related to tolerance [1]. However, few studies have identified central and possible universal regulation of the response to multiple disturbances, a concept termed as perturbome [4,5]. The common molecular response was previously reported as a network of a common set of genes and pathways that can be generically associated with multiple perturbations in plants [7], pathogenic bacteria [3], or cell lines models [4], and others.

In our approach using *P. aeruginosa*, but also applicable to other organisms, we hypothesized the existence of a set of core genes regulating the response to stressors in a generic sense of different pathways. This bacterium is a ubiquitous organism that causes infections in immunocompromised hosts [14]. It has a high proportion (about 5%) of its genome dedicated to regulatory mechanisms, probably explaining its adaptability to such a broad range of growth conditions [38]. Since the strain *P. aeruginosa* PAO1 is a clinical isolate with a profile of multiresistance to many antibiotics [57], characterization of molecular mechanisms involved in the tolerance to stressors in this strain could eventually help to modulate sensitivity and overcome resistance. Exhaustive integration of −omics data and network analysis are required to clarify the molecular mechanisms related to stress conditions and eventually use them for modulating cell response.

For example, we have a particular interest in *P. aeruginosa* due to we have been studying a high-risk strain (called *P. aeruginosa* AG1) which was isolated in 2010 from a Costa Rican Hospital. We reported this isolated in 2015 [58] and it is a Priority 1 organism [59] because it is resistant to carbapenems, a last-resort antibiotic. We assembled the genome using Illumina and Nanopore sequencing [13], revealing the genomic features including two integrons, which were studied in detail in another work [60]. Proteomic profiles after exposure to antibiotics have been also studied [61]. Finally, we studied the transcriptome in response to ciprofloxacin using RNA-Seq and systems biology [16]. We expect to integrate our results of the perturbome (in the reference strain) with the Costa Rican strain (AG1) to identify the “expected general response” to multiple perturbations (as a Pseudomonal), and more importantly, to identify the “exclusive response” to antibiotics, to perform new assays. In the response to ciprofloxacin, we already demonstrated that energy metabolism, RNA degradation, and stress response are induced, as found in other perturbations (that here we call “the perturbome”), but also other more exclusive responses such as phage induction (very exclusive and not part of the perturbome) are modulated with ciprofloxacin.

On the other hand, although nowadays RNA-Seq is one of the most common high-throughput molecular strategies to analyze transcriptomic data, the selection of microarray data to analyze the perturbome of *P. aeruginosa* was based on the availability of data in databases. First, unlike other microorganisms (for example intra-cellular bacteria), *P. aeruginosa* has a very variable genome size between 6 and 7.5 Mb, which is explained in part due to the great repertory of the accessory genome [13]. Because of this, it is difficult to combine molecular data of multiple strains. Second, the PAO1 strain is the reference strain (model for the group during more than 20 years), and no other strain has the completeness of gene annotation as found for this model. Finally, transcriptomic data of PAO1 is mainly found with microarray strategies, which is significantly more abundant in databases than RNA-Seq.

### 4.1 Insights of algorithms to identify core response genes or perturbome

In our study, initial global transcriptomic profiles showed mixed patterns between samples according to their experimental condition. Because of its complexity, raw microarray data showed noise and redundant information that can explain the poor resolution of the initial PCA to identify classes [33]. Thus, a first step was the normalization, which was particularly critical for multi-origin samples as in our case. Comparative analysis needs to consider variation that depends not only on the experimental design but also parameters including laboratory/culture conditions and mRNA extraction methods [62]. Subsequently, a feature (gene) selection analysis was implemented for not only identifying genes associated with multiple perturbations in *P. aeruginosa*, but also to improve the performance of predictive/descriptive models capable of separate control and perturbation categories. In three experiments, two or three time points were considered as perturbations (Figure 3-A and the example of Experiment 3 in Figure 2-B). Although an overrepresentation of that perturbation cannot be discarded, we assume they are independent assays due to the global profile is known to be different over time.

The potential patterns of the core response were investigated using a robust machine learning approach by implementing six protocols, using SP and MP methods, and SVM, KNN, and RF algorithms in each case. This was a critical step because gene selection from microarray data is complex *per se.* Feature reduction remains a challenging task in transcriptomic studies because thousands of genes to select from, and it introduces an additional layer of complexity in the modeling task [12,63]. To avoid bias and overfitting, implementations of diverse strategies of data partitioning such as bootstrapping, random partitions, and cross-validation are recommended. In general, these methodologies can robustly minorate the influence from noise, outliers, absence of ground truth sets, and reduce variance [2,37,64]. In the specific case of implementations for the study of molecular networks, partitions are suggested not only because to counteracts the curse of dimensionality but also ground truth of candidate genes or regulatory networks are rarely available to allow for an unbiased assessment of these methods [37].

The single partition SP method consisted of particular and invariable data for training (with internal cross-validation) and another to test, and it is probably the most common approach used in machine learning. In the case of the multiple partitions MP method, 100 random partitions of the dataset were run. MP method had a dual consideration when splitting data (multiple partitions and the internal cross-validation). This method can be considered as an *ensemble based on different data partitioning*, as it had been previously proposed [36]. Datasets were divided using multiple random partitioning procedures and then genes were ranked. After all runs, a final feature subset is determined by calculating the frequency of features in all the runs [36]. An equivalent approach was implemented by Pai and collaborators to classify gene expression data in a cancer model [65].

However, one possible problem with the MP approach is that cross-validation results may depend on the similarity of testing and training sets. For example, all the samples of the experiment can be assigned as part of the same training set and none in the testing set (or vice versa, for example, the red/blue/dark-green replicates of the last arrangement of the MP method in Figure 2-B). A classification prediction method is only expected to learn how to predict on unseen samples that are drawn from the same distribution as training samples [37,64], and the MP method could violate this assumption. SP method guarantees this because it was built using the last replicate for each perturbation and control (Figure 2-B, SP method).

In contrast, because of the presence of several biological replicates of the same perturbation in both datasets of the SP method (one in the testing and the remaining in the training set), it can be considered that the conformation of the testing data set is not completely independent, i.e. every sample in the training dataset has a biological “copy” (replicate) itself in the training set. This scenario can introduce a bias in the performance of the classifiers and robust analyses, such as multiple partitions and/or cross-validation, are required to avoid a false impression of high predictive ability of the model and hence of the reliability of the inferred influences on the gene [37].

Thus, both SP and MP methods have drawbacks but they can be compensated using the best features of each strategy to robustly select features (genes).

On the other hand, many algorithms for dimension reduction have been proposed [6,32,33] but no standard machine learning algorithm can be selected due to multiple evaluation results on completely variable metrics associated with the performance [39]. Many studies have shown that SVM, KNN, and RF generally outperform other traditional supervised classifiers [30,39,66]. In our case, a variable pattern was found for different metrics in all evaluated implementations. For example, based on accuracy SVM for SP method and RF for MP method resulted in higher scores, which agree with other studies using machine learning and biological data [39,65].

In our subsequent analysis, we were interested in identifying a consensus list of candidate genes for the core response in *P. aeruginosa*, resulting in 46 genes of the perturbome. When global profiles were compared using these genes (PCA), control and perturbation classes were clearly separated (Figure 7-C). This gene number seems to be a modest number of elements (less than 1% of all the available datasets with 5549 genes) but it agrees with other studies, including machine learning methods [11,30] or other approaches [11,35,67,68]. Besides, 72% of core response genes were also identified as DEGs with a similar annotation profile. Differences can be explained by significant fluctuations in the differential expression results as previously reported, mainly because it is not a consensus strategy (only based on *p*-value) and it does not incorporate the estimates of the test performance (true positive/negative rates and other metrics) on the results [2].

### 4.2 Biological insights of the core response genes in *P. aeruginosa*: the perturbome

In the study of complex systems, the lack of specific biological information at the single gene level and the interactions make the extraction of the relevant information a complicated task in this field. This situation remains as an unsolved issue of molecular high-throughput technologies, in which molecular data is generated at a higher rate than our ability to find patterns or extract relevant information [69]. Thus, large-scale/top-down system biology-based approaches have been used as a strategy to exploit and extract relevant information of ample amount of −omics data [21], letting to predict interactions between genes [23].

With the representation of gene-to-gene interactions, system biology allows studying not only specific genes or gene clusters but also the emergent properties found in the molecular networks [17]. Thus, pleiotropic patterns can be recognized towards the response to perturbations, in which large-scale networks can be built [69]. In the case of the core perturbome, the representation of genes showed a complex network with topological properties that defines a non-random arrangement, including hub genes and pathways associated with the response to stress and stimulus, metabolism, and ribosome activity. The network can be inferred as a coordinated central response to the perturbations, independently of a particular stressor.

Genes of the core response or perturbome can be related to a central regulation network, and as the convergent point of signal transduction, transcriptional regulation, and stress-related pathways, as it has been suggested [2,4,5,56]. Annotation of the 46 genes shows that most of them are functionally related to biosynthesis, molecule binding, and metabolism (including an important number of hits for lipids), including additional functions for the regulation of DNA damage repair, response to stimuli, and aerobic respiration. Interestingly, hub genes represent these processes with high connectivity in the small-world network. For example, the main functions associated with *fabA* are lipid metabolism, fatty acid, and lipid biosynthesis, meanwhile for *nuoC* are aerobic respiration, electron/proton transport, and DNA damage. Finally, *nuoF* is associated with aerobic respiration, electron/proton transport, and Krebs cycle.

In this sense, cells are equipped with systems and mechanisms to recover from the environmental stress and stimuli to maintain all necessary physiological functions [70]. Other common stimuli such as low ATP, slow growth, and ROS production can also occur before cells express stress-specific factors, but mediating a common effect. For example, response to stress includes modulation of energy production and aerobic electron transfer chain components. As it has been reported in *E. coli*, aerobic electron transport chain components are down-regulated in response to growth arrest [71]. This corresponds with the global profiles of expression of 46 core response genes. Also, regulation of lipid metabolism is relevant for survival in the wide range of environmental conditions where bacteria thrive [72], even for biofilm-living forms [3] as *P. aeruginosa.* Core response genes *fabG, lpxA*, and PA5174 could be implied in this process.

In the case of DNA damage repair (including the case of *cycB* and *gltP* genes), responses mediated by SOS and *rpoS* help to maintain genome integrity, colonization, and virulence [19,73]. These responses are activated under multiple disturbances and modulating a low energy production and shutdown of the metabolism, promotes the formation of antibiotic resistance and biofilms [3]. Other related pathways for some specific genes included regulation of the transcription during stress by RNA-binding proteins to reprogram or shut down translation and to rescue the ribosome stalled by a variety of mechanisms induced [74]. Three core response genes (*rpmH, tsf*, and PA2735) were annotated with these functions.

Jointly, the relatively few diversity of metabolic functions and pathways makes sense in order to ensure redundancy and robustness in the response to stimuli. Similar results, regarding enriched pathways, have been obtained in other studies with eukaryotic models, including disturbed human cell lines [4,5,56], *Arabidopsis thaliana* under physical and genotoxic stresses [2], or a genome-wide association study of a generic response to stress conditions [6]. In the case of prokaryotic organisms, two studies have used *Escherichia coli* as a model to identify differentially expressed genes after exposure to stress conditions [3] and to create networks associated with the response to fluctuating environments [7]. Differences with other organisms and disturbances can suggest that response cell stress can be organismal specific, although heterogeneity has also been suggested as a reasonable explanation because of differences in the response in a homogeneous cell population [56,75].

As in our case, the response to stimuli and stressors is orchestrated by a pleiotropic modulation [20] which can be associated with central regulation. Alternative mechanisms such as cross stress protection (the ability of a stress condition to protect against other stressors) [7], the role of sigma factors, and specific two-component systems [3] can contribute to explaining this phenomenon. The molecular response can lead to regulate multiple biological activities including metabolism, replication, transcription and translation, changes in membrane composition, motility, modification gene expression, expression of virulence factors, multi-drug resistant phenotypes and biofilm formation, and others [56].

For further works, we are running experiments to validate our findings here, in which *P. aeruginosa* is exposed to different perturbations and gene expression is quantified by RT-qPCR. This will also allow us to build small-world networks with the representation of the core perturbome as a directed graph. Using a bottom-up system biology approach using differential equations, sensitivity analysis, and other machine learning algorithms, we expect to identify and manipulate the points of control the network, as we have previously done in other models [76].

Taking all together, results of our study suggest that identification of core response genes associated with multiple perturbations or perturbome in *P. aeruginosa* can define a central network available to modulate a basic response that includes biological functions such as biosynthesis, binding, and metabolism, many of them related to DNA damage repair and aerobic respiration. To our knowledge, this study can be considered a first report of the *P. aeruginosa* perturbome.

Further analyses are required to explore the potential use of perturbome network to modulate (positively or negatively) the response to disturbances, to model molecular circuits, to identify possible biomarker genes of stressed states, and to experimentally validate our findings. Besides, this approach can be used to model the perturbome in other *P. aeruginosa* strains, as we hope to run soon with a genome we recently described [13], and other organisms.

## 5. Conclusions

A robust machine learning approach was implemented to identify and describe core response genes to multiple perturbations in *P. aeruginosa.* Using public microarray data, two independent partition strategies (single and multiple with SP and MP methods respectively), and three classification algorithms, we were able to identify 46 perturbome elements. Both network analysis and functional annotations of these genes showed coordinated modulation of biological processes in response to multiple perturbations, including metabolism, biosynthesis, and molecule binding, associated with DNA damage repairing, and aerobic respiration, all probably related to tolerance to stressors, growth arrest, and molecular regulation. We also discussed different issues related to implemented and assess algorithms of normalization analysis, data partitioning, classification approaches, and metrics.

## Abbreviations

AUC: Area under the curve
AZM: Azithromycin
B8F: E-4-bromo-5-bromomethylene-3-methylfuran-2-5H-one
CDA: Chlorhexidine diacetate
CIP: Ciprofloxacin
COL: Colistin
Cu: Copper
DEGs: Differentially expressed genes
GEO: Gene Expression Omnibus
H2O2: Hydrogen peroxide
HC: Hierarchical Clustering
KNN: K-Nearest Neighbor
LB: Luria Bertani
mRNA: Messenger RNA
NaClO: Sodium hypochlorite
OPP: Ortho-phenylphenol
PCA: Principal Component Analysis
PPI: Protein-protein interaction
RF: Random Forest
ROC: Receiver-operating characteristic
SVM: Support Vector Machine

## Declarations

### Ethics approval and consent to participate

Not Applicable

### Consent for Publication

Not Applicable

### Availability of data and material

The datasets generated and/or analyzed during the current study are available in: Public raw data used in this study: GEO database (https://www.ncbi.nlm.nih.gov/geo/), data Series GSE2430, GSE3090, GSE4152, GSE5443, GSE7402, GSE10605, GSE12738, GSE13252, GSE14253 and GSE36753.

Normalized data and R Scripts: https://github.com/josemolina6/CoreResponsePae

### Authors’ contributions

JMM and FGS participated in the conception, design of the study, and data acquisition. JMM, PMM, RGB, and JVF participated in data analysis and interpretation. JMM, RCS, JVF, and FGS participated in the interpretation of the data analysis. JMM drafted the manuscript and all authors were involved in its revision. All authors read and approved the final manuscript.

## Acknowledgments

We thank students Iosif Forero Trelles, Daniel Solano Alvarado, and Daniela Aguilar Orozco for their collaborations in different activities of the project.

## Supplementary Figures and Tables

**Figure S1.**
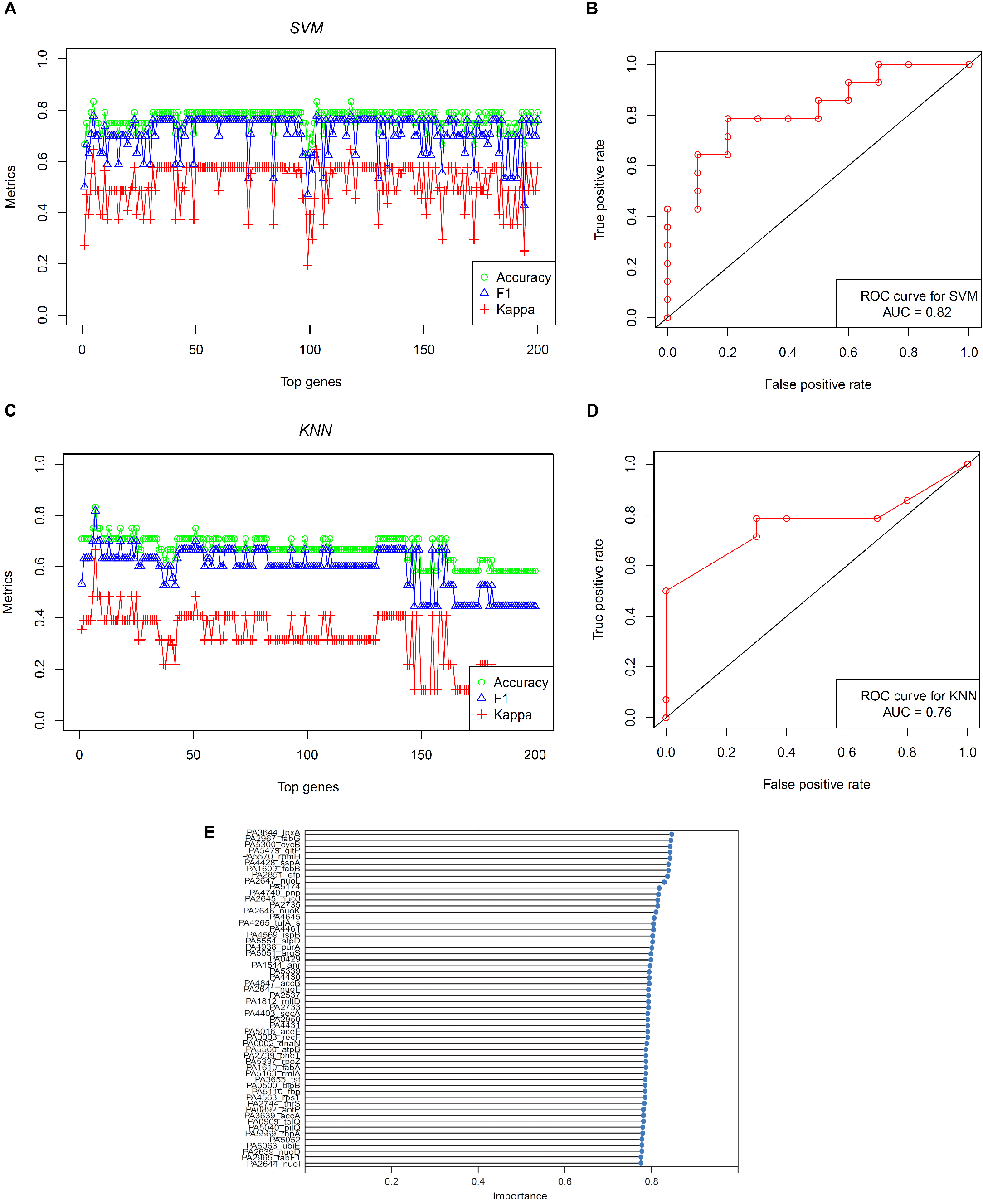
Evaluation of SP method for gene ranking by importance, the case for SVM and KNN algorithms. (A-C) Accuracy, F1, and kappa values after iterations of classification with the first top 200 genes (adding genes 1-to-1). (B-D) ROC plot using selected top 56 genes for evaluation of the performance of the algorithm. (E) Ranking and importance value of top 56 genes (same importance for both algorithms).

**Figure S2.**
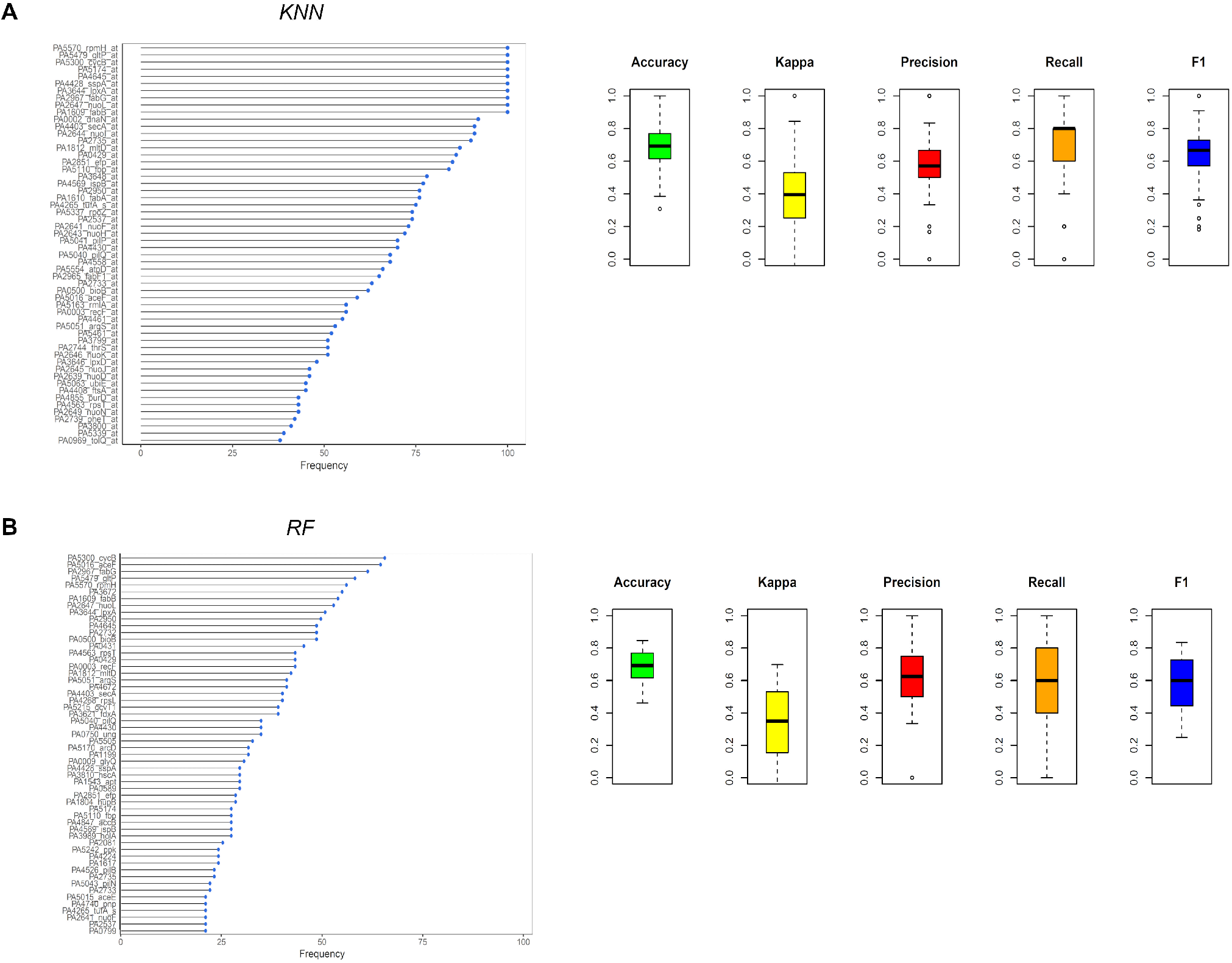
Evaluation of MP method for gene ranking by frequency, the case for KNN and RF algorithms. (A) KNN evaluation of ranking of top 56 genes by frequency after iterations of 100 data partitions and classification model fitting and dispersion of metrics across 100 iterations. (B) Same evaluation, the case for RF algorithm.

**Figure S3.**
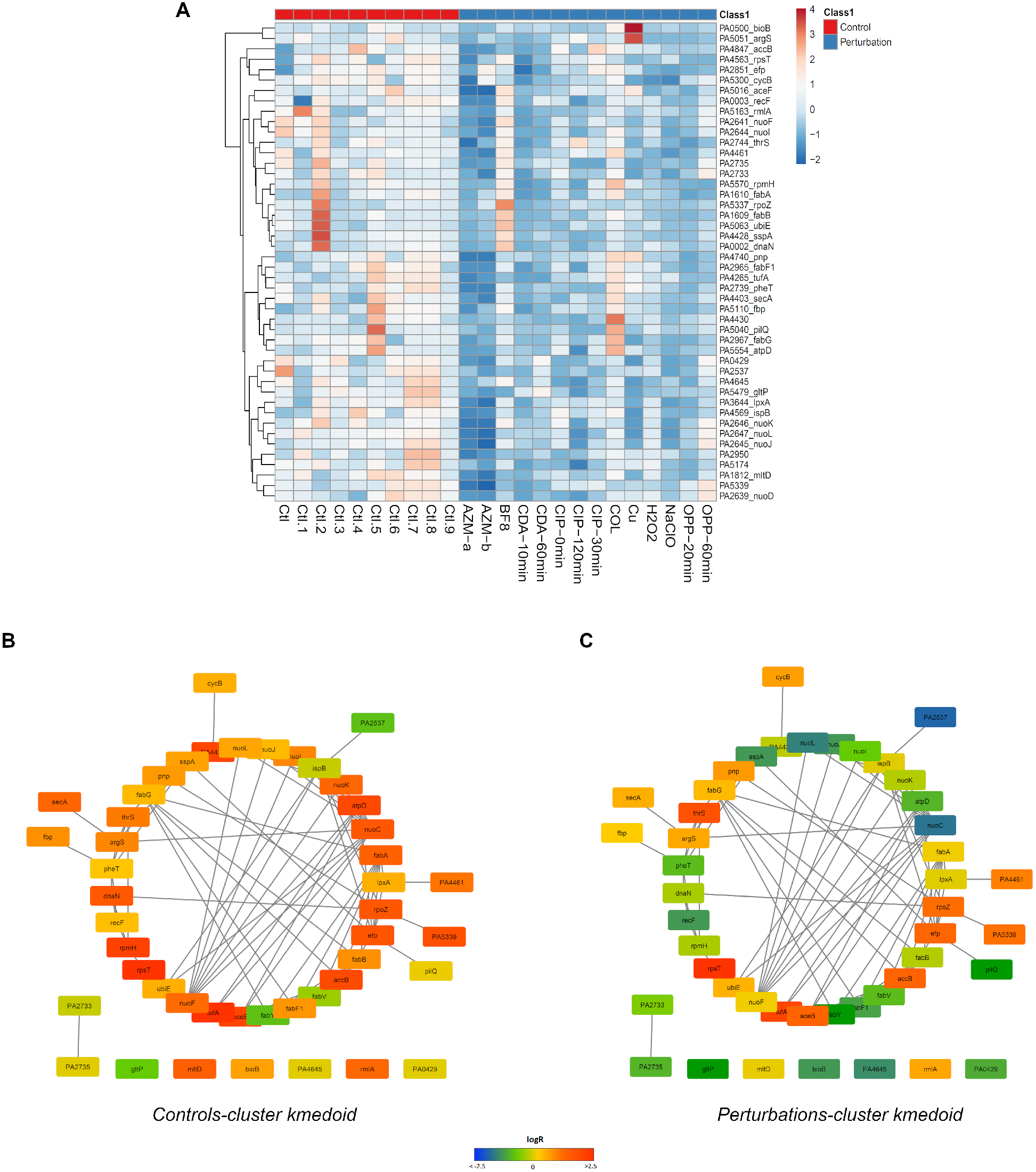
Systems-level description and gene expression comparison by core response genes. (A) Gene expression and clustering of the 46 core response determinants of *P. aeruginosa.* (B-C) Comparison of gene expression (logR) of kmedoids for each cluster using small-world networks.

**Table S1.**
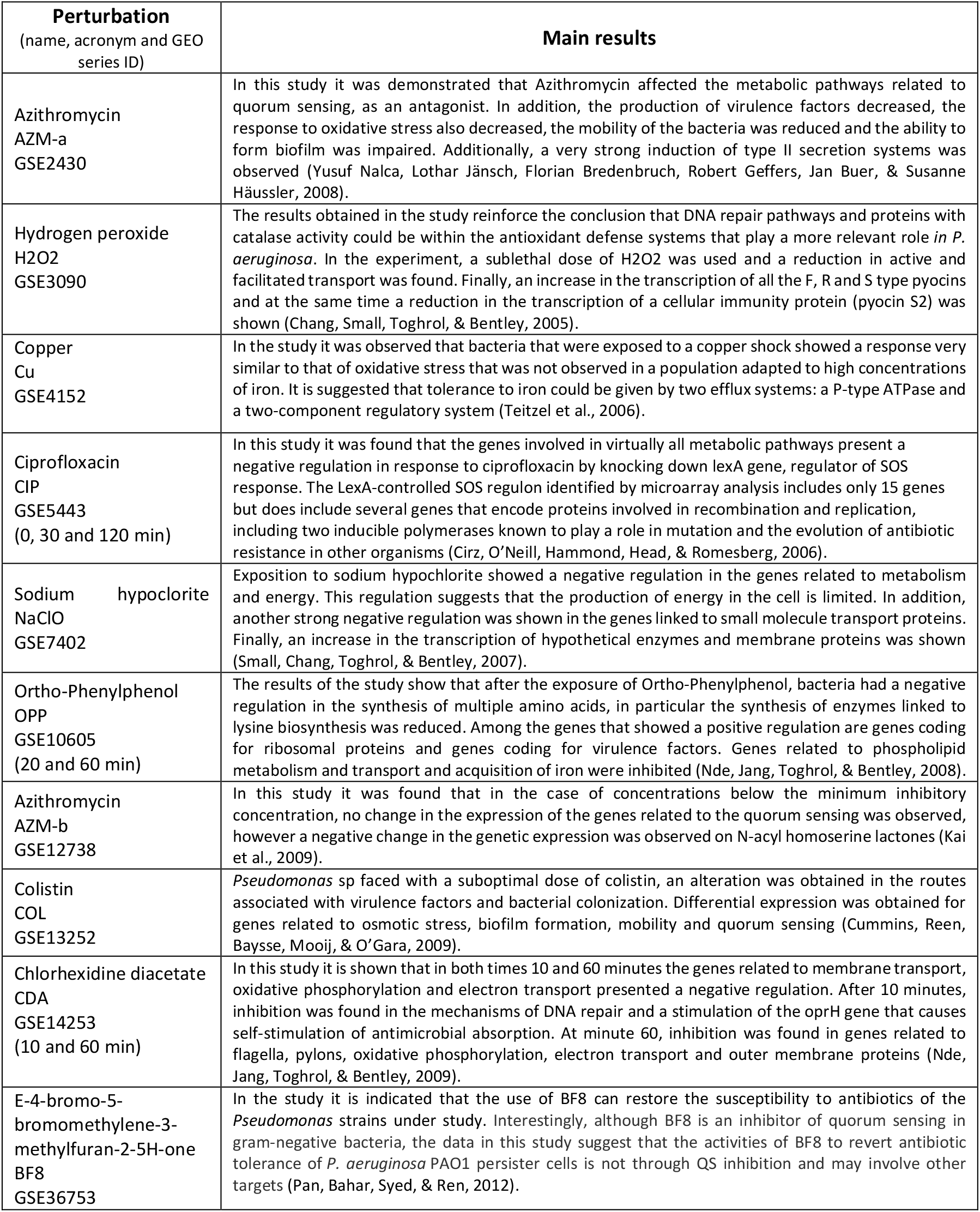
Studies of datasets included in this work to identify the core perturbome of *Pseudomonas aeruginosa*

**Table S2.**
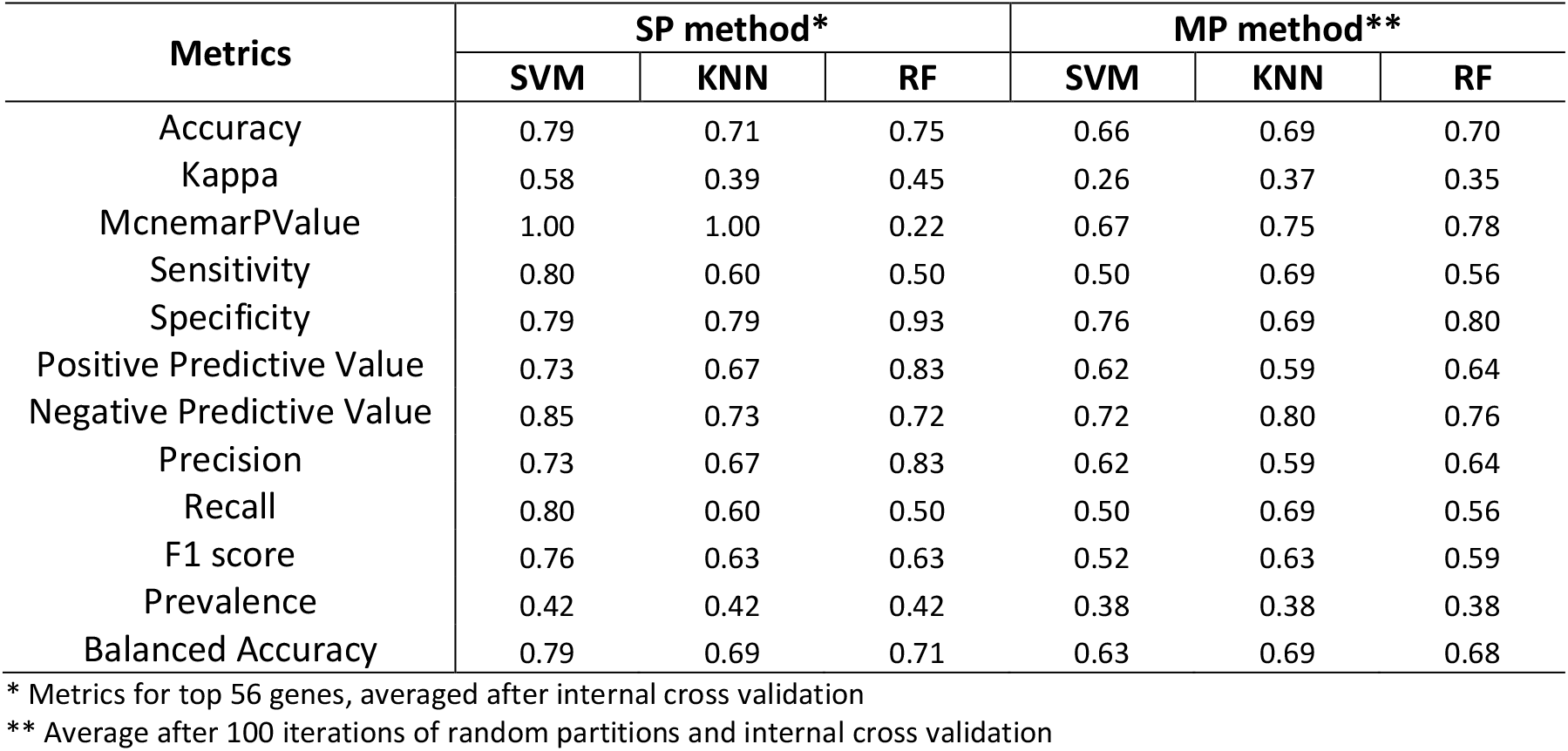
Metrics associated with the performance of each method and algorithm

